# Cortical tethering of mitochondria by the dynein anchor Mcp5 enables uniparental mitochondrial inheritance during fission yeast meiosis

**DOI:** 10.1101/525196

**Authors:** Kritika Mehta, Vaishnavi Ananthanarayanan

## Abstract

During sexual reproduction in eukaryotes, processes such as active degradation and dilution of paternal mitochondria ensure maternal mitochondrial inheritance. In the isogamous organism fission yeast, we employed high-resolution fluorescence microscopy to visualize mitochondrial inheritance during meiosis by differentially labeling mitochondria of the two parental cells. Remarkably, mitochondria, and thereby, mitochondrial DNA from the parental cells did not mix upon zygote formation, but remained segregated at the poles by attaching to clusters of the dynein anchor Mcp5 via its coiled-coil domain. We observed that this tethering of parental mitochondria to the poles results in uniparental inheritance of mitochondria, wherein two of the four spores formed subsequently contained mitochondria from one parent and the other spores, mitochondria from the other parent. Further, the presence of dynein on an Mcp5 cluster precluded the attachment of mitochondria to the same cluster. Taken together, we reveal a distinct mechanism that achieves uniparental inheritance by segregation of parental mitochondria.

## Introduction

Mitochondria are cellular organelles responsible for the generation of energy-rich adenosine triphosphate molecules in eukaryotic cells. In addition to this and other essential functions, mitochondria carry their own genetic material in the form of mitochondrial DNA (mtDNA) nucleoids. During meiosis, in contrast to the nuclear genome, mitochondrial genes follow a non-Mendelian pattern of segregation through tightly controlled mechanisms that typically favor uniparental inheritance, or the passing down of mitochondria predominantly from a single parent to the progeny. In several eukaryotes, maternal inheritance is the preferred mode of uniparental inheritance. Maternal inheritance is brought about by one of many ways including subjecting paternal mitochondria to (i) sequestration and exclusion (Yu and Russell, 1992), (ii) selective lysosomal degradation via ubiquitination (Sutovsky et al., 1999, 2000), (iii) simple dilution due to the large size of the female gamete in comparison to the male gamete (Birky, 1995; Wilson and Xu, 2012). Uniparental mitochondrial inheritance has been suggested to be important for preventing the propagation of selfish cytoplasmic transposable elements that could affect the nuclear genome (Hoekstra, 2000; Murlas Cosmides and Tooby, 1981).

In the unicellular eukaryote budding yeast *Saccharomyces cerevisiae*, mitochondria are biparentally inherited by the meiotic progeny due to mixing of mitochondria from both parental cells upon zygote formation (Strausberg and Perlman, 1978; Thomas and Wilkie, 1968; Zinn et al., 1987). However, mitochondrial DNA (mtDNA) that occur in the form of nucleoids seemingly remain anchored to their original locations in the zygote, thereby giving rise to a homoplasmic cells within a few rounds of vegetative division following sporulation (Nunnari et al., 1997). During mitosis in *S. cerevisiae*, mitochondria in the mother cell are tethered to the cell membrane via the mitochondria-ER cortex anchor (MECA) structure containing the protein Num1 (Lackner et al., 2013; Ping et al., 2016). Tethering of mitochondria by Num1 aids in the retention of a mitochondrial population within the mother cell (Lackner et al., 2013), while another population is transported on actin cables to the bud by the activity of the myosin V, Myo2 (Altmann et al., 2008; Förtsch et al., 2011). The Num1 homologue in fission yeast (*Schizosaccharomyces pombe*), Mcp5 is expressed specifically during prophase I of meiosis (Saito et al., 2006; Yamashita and Yamamoto, 2006), and is required for the anchoring and thereby, activation of the motor protein cytoplasmic dynein that powers the oscillatory movement of the zygotic horsetail-shaped nucleus (Ananthanarayanan et al., 2013; Tolic et al., 2009; Yamamoto et al., 1999).

Interphase mitochondria in fission yeast remain associated with microtubules and their fission dynamics are dictated by the dynamics of the underlying microtubules (Chiron et al., 2008; Fu et al., 2011; Yaffe et al., 1996). This relationship between microtubules and mitochondria is also essential for independent segregation of mitochondria during mitosis (Mehta et al., 2019). However, it is unclear how mitochondria are segregated between the four spores that result from meiotic cell division in fission yeast. It has been suggested that like *S. cerevisiase, S. pombe* also undergoes biparental mitochondrial inheritance in crosses between strains resistant and sensitive to antibiotics (Thrailkill et al., 1980), but direct evidence for this process in wild-type cells has been lacking.

Here, we report that fission yeast cells in fact undergo uniparental mitochondrial inheritance during meiosis due to the tethering of mitochondria to the cortex during the initial stages of meiosis. Our results thus reveal a unique mechanism for facilitating uniparental inheritance that relies on physical segregation of parental mitochondria in a heteroplasmic zygote by the activity of the dynein anchor Mcp5.

## Results

### Mitochondria are preferentially localized at the poles of meiotic cells

To the best of our knowledge, there exists no comprehensive study on the changes of the mitochondrial network upon onset of meiosis in fission yeast. Therefore, we first set out to visualize mitochondria during the fission yeast meiotic cycle. We achieved this by inducing meiosis in parental cells that had fluorescently labeled mitochondria and microtubules (Figure 1A, Movie S1), or mitochondria and nucleus (Figure 1B, Movie S2).

**Figure 1.**
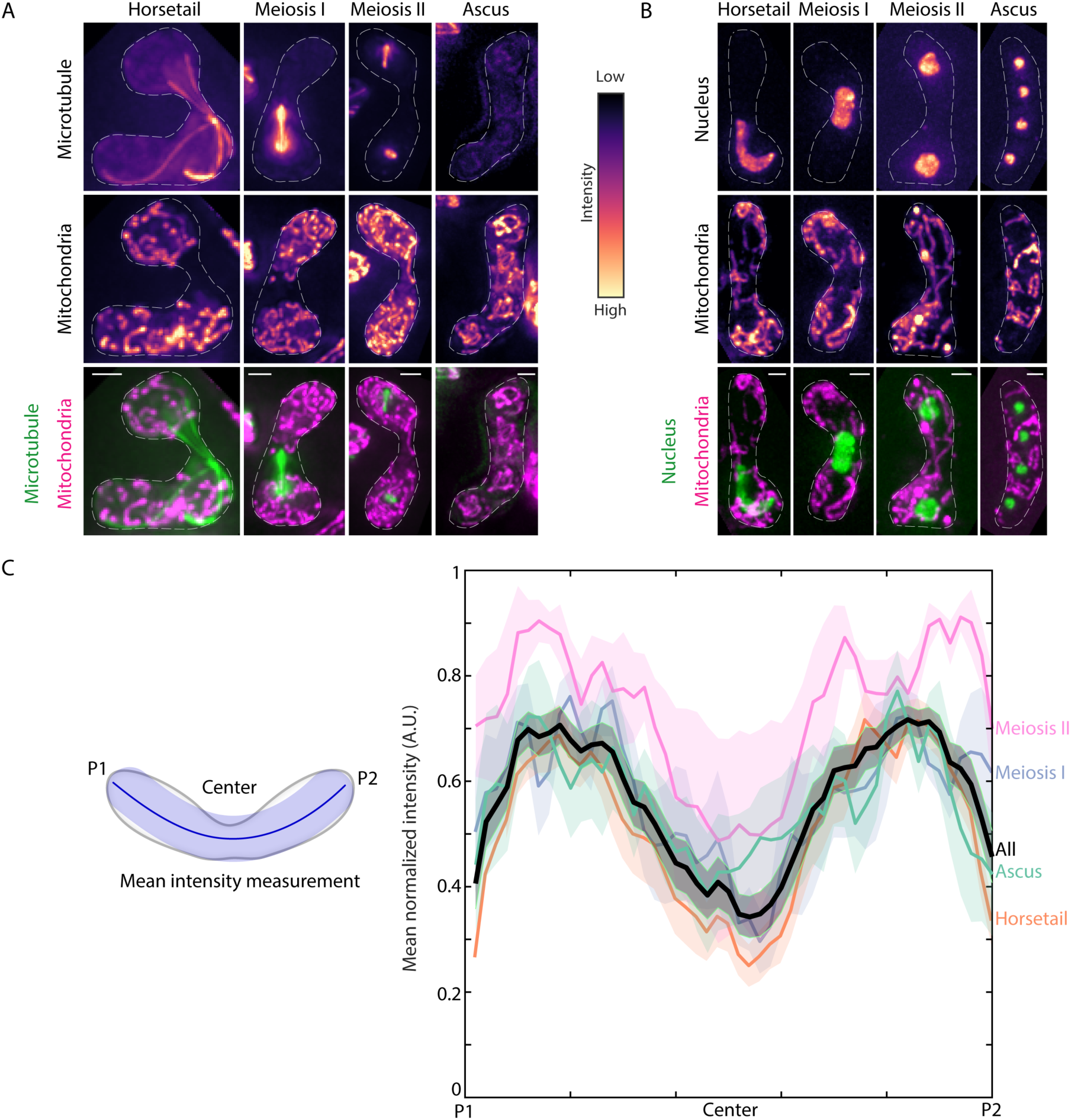
Mitochondria remain close to the cell poles during meiosis. **(A)** Maximum intensity-projected images of microtubules (top) and mitochondria (middle) represented in the intensity map to the right of the images, and their merge (bottom) during the different stages of meiosis indicated (strain KI001xPT1651, see Table S1). **(B)** Maximum intensity-projected images of the nucleus (top) and mitochondria (middle) represented in the intensity map to the left of the images, and their merge (bottom) during the different stages of meiosis indicated (strain FY15112, see Table S1). In A and B, scale bars represent 2µm, dashed white lines represent cell outlines. **(C)** Schematic (left) of the mean intensity measurement along the length of a zygote from pole ‘P1’, through the ‘Center’, to pole ‘P2’. Plot of mean normalized intensities (right) from different stages of meiosis (colored lines) and their combined mean intensities (black line, *n*=24) obtained from the data in A. The shaded regions represent the standard error of the mean (SEM).

Based on the microtubule organization and the nuclear morphology, the discernible stages of meiosis were designated as ‘horsetail’, ‘meiosis I’, ‘meiosis II’ and ‘ascus’ (Cipak et al., 2014). In contrast to interphase mitochondria, during meiosis, mitochondria appeared predominantly fragmented and detached from the microtubules (e.g.: Figure 1A, ‘horsetail’). Further, the mean normalized intensity of mitochondria across the cell for all stages revealed preferential localization of mitochondria to the poles of the cell (Figure 1C).

### Parental mitochondria do not mix upon zygote formation

Next, we sought to understand how mitochondria are inherited during fission yeast meiosis. To this end, we employed cells of opposite mating types whose mitochondria were labelled with different fluorophores, GFP and RFP. We induced meiosis in these cells and observed mitochondrial organization during the early horsetail stage and in the final stage, post formation of ascospores. Interestingly, we observed that the differently labelled mitochondria from the parental cells remained segregated at the poles of the cell and did not undergo any mixing in the early stage (Figure 2A, top, Movie S3). Upon formation of spores within the ascus, mitochondria remained predominantly unmixed, with two of the spores exhibiting higher GFP signal and the two other, higher RFP signal (Figure 2A, bottom, Movie S4). These observations were consistent with our measurement of normalized mean mitochondrial intensities across the length of the cell (Figure 2B). We additionally visualized meiotic mitochondrial inheritance in a cross between a cell containing fluorescently-labeled mitochondria and a cell containing unlabeled mitochondria. Here too, we observed localization of mitochondrial signal to one side of the zygote and two spores of the resulting ascus (Figure S1A, B).

**Figure 2:**
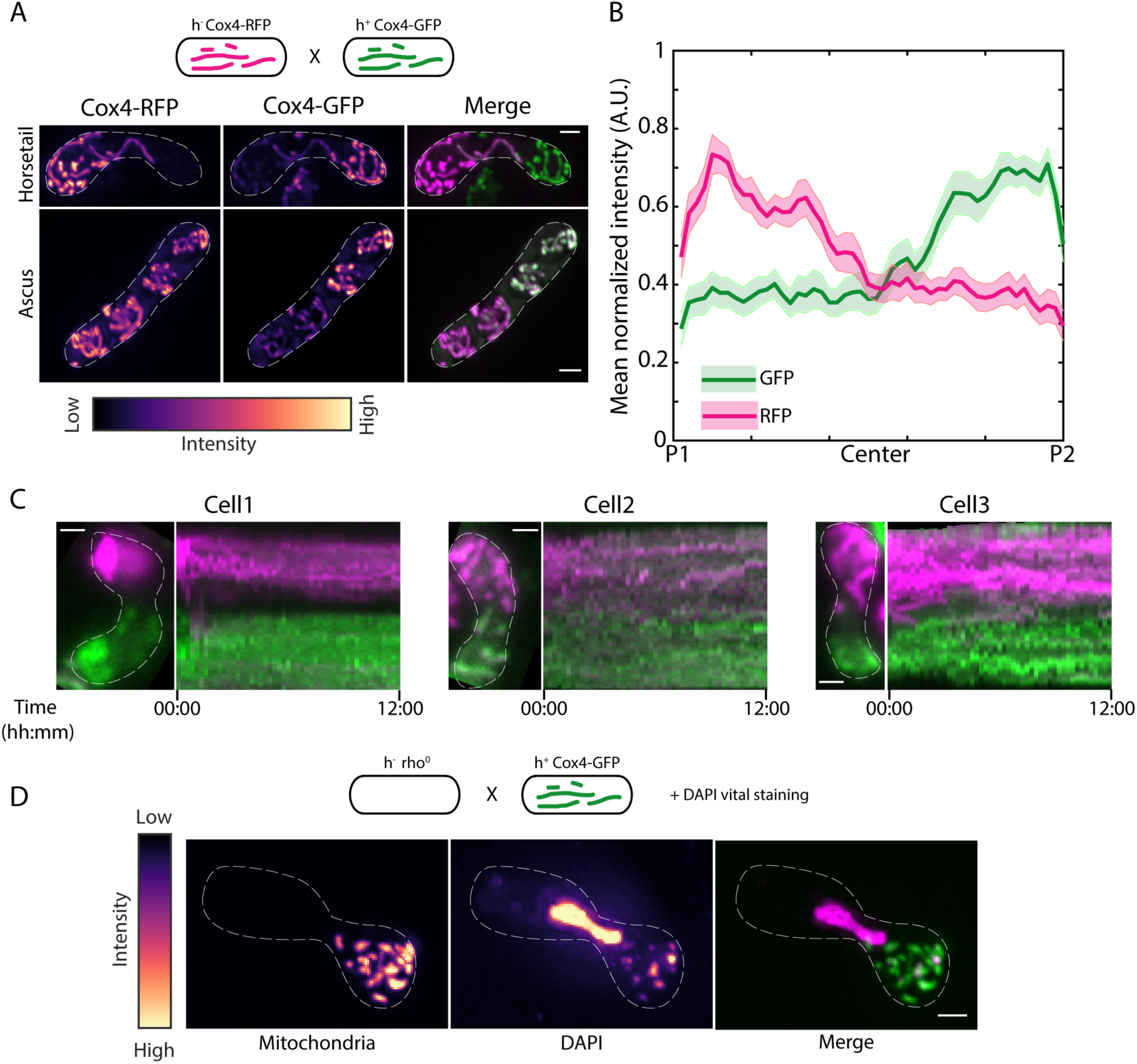
Parental mitochondria remain segregated upon conjugation. **(A)** Schematic of the cross performed (top, strain PT1650xPT1651, see Table S1), maximum intensity-projected images of mitochondria labeled with Cox4-RFP (left) and Cox4-GFP (center) represented in the intensity map to the bottom of the images, and their merge (right) during the early stage (‘Horsetail’, top) and late stage (‘Ascus’, bottom) of meiosis. **(B)** Plot of mean normalized intensities of RFP (magenta line) and GFP (green line) across the length of the cell from the cross indicated in A (*n*=18). Shaded regions represent SEM. **(C)** Representative maximum intensity-projected images (left) and kymographs of time-lapse movies (right) of meiotic cells resulting from the cross indicated in A. Time is indicated below the images in hh:mm. **(D)** Schematic of the cross and DAPI vital staining performed (top, strain PHP14xPT1650, see Table S1), maximum intensity-projected images of mitochondria labeled with Cox4-GFP (left) and mtDNA (‘DAPI’, center) represented in the intensity map to the left of the images, and their merge (right). In A, C and D, scale bars represent 2µm, dashed white lines represent cell outlines.

In all these experiments, the mitochondrial inner membrane protein Cox4 was used as a fluorescent reporter for the mitochondria. To rule out any effects from differential dynamics of the mitochondrial compartments (Sukhorukov et al., 2010), we employed another fluorescent reporter protein for the mitochondrion that resides in the mitochondrial matrix, aconitase (Aco1) tagged with GFP. Again, we observed segregation of the mitochondria in meiotic cells resulting from a cross between cells with unlabeled mitochondria and cells with mitochondria labeled with Aco1-GFP (Figure S1C, D).

The segregation of mitochondria that we observed could result from a scenario where mitochondria underwent mixing upon zygote formation, but then subsequently de-mixed via a different process. To test if this occurred, we acquired long-term time-lapse videos of fission yeast cells undergoing meiosis. Again, we employed parental cells with differently labeled mitochondria. We observed that the segregation of mitochondria occurred very early in the meiotic cycle and was maintained during the later stages (*n* = 13, Figure 2C, Movie S5).

To confirm that the segregation of parental mitochondria resulted in segregation of the parental mitochondrial DNA (mtDNA), we first set up a cross between a cell expressing fluorescently labeled mitochondria and a cell lacking mtDNA nucleoids (rho^0^) (Haffter and Fox, 1992). Then, we labeled mtDNA by vital 4’,6-Diamidino-2-phenylindole dihydrochloride (DAPI) staining of the resulting zygote (Williamson and Fennell, 1979). In such a scenario, all the mtDNA in the products of this cross would originate from the non-rho^0^ parental cell. Accordingly, we again observed mitochondrial segregation in the zygote and also observed complete colocalization between the labeled mitochondria and mtDNA (n = 11, Figure 2D, Movie S6). These results indicate that mitochondria and hence mtDNA of the parental cells remain segregated during meiosis and are thereby uniparentally inherited in the meiotic progeny.

### The dynein anchor Mcp5 tethers mitochondria to the poles during prophase I of meiosis

In budding yeast, the Mcp5 homologue Num1 forms a part of the MECA structure and is essential for retention of mitochondria in the mother cell while the Myo2 motor carries mitochondria to the bud on actin cables (Lackner et al., 2013). The mitochondrial localization at the poles that we observed (Fig. 1 and Fig. 2A) was reminiscent of the organization of Mcp5 spots at the cortex (Saito et al., 2006; Thankachan et al., 2017; Yamashita and Yamamoto, 2006). Mcp5 clusters into about 30 foci containing ∼10 molecules per focus, preferentially at the cell poles (Thankachan et al., 2017). Additionally, Mcp5 is a meiosis-specific protein that is expressed predominantly during meiotic prophase in fission yeast, when it anchors dynein to enable oscillations of the horsetail nucleus (Saito et al., 2006; Yamashita and Yamamoto, 2006).

Therefore, to test if mitochondria were also being anchored by Mcp5 in fission yeast, we first visualized zygotes which expressed fluorescently labelled mitochondria and Mcp5. We observed complete colocalization between mitochondria at the cortex and Mcp5 foci (Figure 3A). In this cross, GFP-labelled Mcp5 was expressed from only one of the parents and RFP-labeled Cox4 was expressed from the other. Interestingly, while Mcp5’s signal was visible at both poles of the cell, mitochondrial signal was again restricted to one pole (Figure 3B, Movie S7, Figures. S2A, S2B), indicating that there were no barriers to diffusion or mixing of other proteins in the zygote. Additionally, mitochondria continued to remain dissociated from the microtubules when bound to Mcp5 (Figure S2C), as observed in Figure 1A. To verify that the attachment to microtubules was not necessary for segregation during meiosis, we employed parental cells lacking the microtubule-mitochondrial linker protein Mmb1 (Fu et al., 2011). Additionally, one of the parental cells had its mitochondria fluorescently labeled. In zygotes and asci resulting from this cross, we observed that parental mitochondria continued to remain segregated (Figures S2D, S2E).

**Figure 3:**
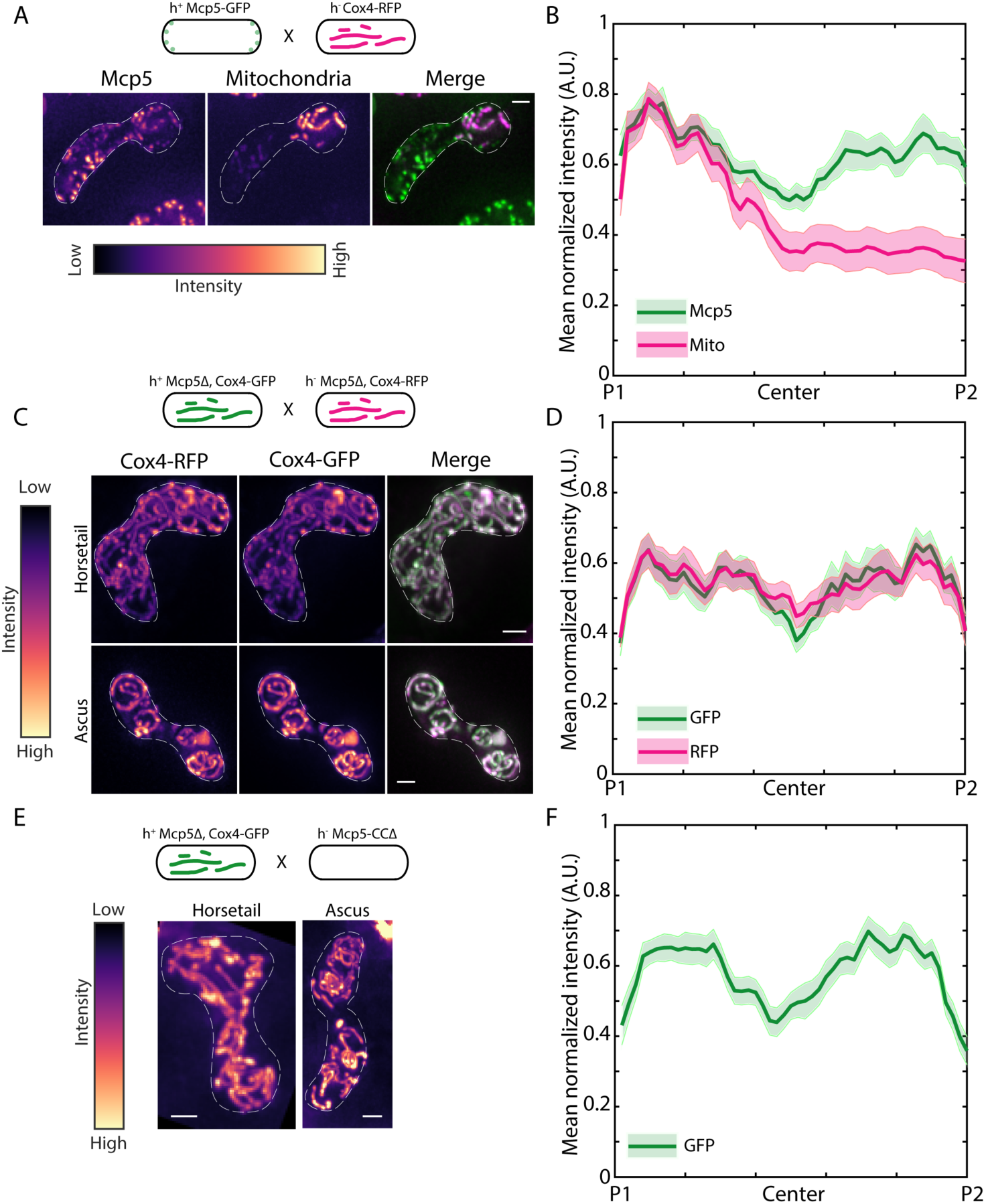
Mcp5 is essential for mitochondrial tethering to the cortex. **(A)** Schematic of the cross performed (top, strain FY16854xPT1651, see Table S1), maximum intensity-projected images of Mcp5 labeled with GFP (left) and mitochondria labeled with Cox4-RFP (center) represented in the intensity map to the bottom of the images, and their merge (right). **(B)** Plot of mean normalized intensities of Mcp5 (green line) and mitochondria (magenta line) across the length of the cell from the cross indicated in A (*n*=14). **(C)** Schematic of the cross performed (top, strain VA066xVA074, see Table S1), maximum intensity-projected images of mitochondria labeled with Cox4-RFP (left) and mitochondria labeled with Cox4-GFP (center) represented in the intensity map to the left of the images, and their merge (right) during the early stage (‘Horsetail’, top) and late stage (‘Ascus’, bottom) of meiosis. **(D)** Plot of mean normalized intensities of RFP (magenta line) and GFP (green line) across the length of the cell from the cross indicated in C (*n*=18). **(E)** Schematic of the cross performed (top, strain FY16897xVA074, see Table S1), maximum intensity-projected images of mitochondria labeled with Cox4-GFP during the early stage (‘Horsetail’, left) and late stage (‘Ascus’, right) of meiosis represented in the intensity map to the left of the images. **(F)** Plot of mean normalized intensity of GFP (green line) across the length of the cell from the cross indicated in E (*n*=21). In A, C, and E, scale bars represent 2µm, dashed white lines represent cell outlines. In B, D and F, shaded regions represent SEM.

We then proceeded to set up a cross between cells lacking Mcp5, but with GFP- and RFP-labeled mitochondria. In stark contrast to wild-type zygotes, these Mcp5Δ meiotic cells showed complete mixing of parental mitochondria in both early and late stages (Figure 3C, Movie S8). These observations were also substantiated by measurement of GFP and RFP intensities across the length of the cell during all stages of meiosis (Figure 3D, Movie S9).

Mcp5 comprises of a pleckstrin-homology domain, which is essential for its attachment to the membrane, and a coiled-coil (CC) domain, that is required for its binding to dynein (Ananthanarayanan, 2016; Saito et al., 2006; Yamashita and Yamamoto, 2006). We asked if the CC domain was also responsible for Mcp5’s attachment to the mitochondria. To answer this, we visualized mitochondrial distribution in a cross between a parental cell lacking Mcp5’s CC domain and the other parent with the genotype Mcp5Δ, Cox4-GFP. If mitochondrial tethering by Mcp5-CCΔ was intact, we would observe an intensity pattern similar to that in Figure S1A or S2A. However, we saw that the fluorescence from the mitochondria was distributed throughout the cell in both early and late stages (Figure 3E and F, Movies S10 and S11), indicating that Mcp5 indeed employs its CC domain to tether mitochondria to the cortex during meiotic prophase I.

### Dynein-Mcp5 spots on the membrane are devoid of mitochondria

When deleting Mcp5 to test its role in mitochondrial tethering, we not only knocked down Mcp5, but also abrogated the oscillations that occur during the meiotic prophase (Saito et al., 2006; Thankachan et al., 2017; Yamashita and Yamamoto, 2006). To delineate the specific role of the oscillations, if any, in facilitating parental mitochondrial segregation, we sought to attenuate the oscillations of the horsetail nucleus while keeping Mcp5 intact. To this end, we employed cells lacking the motor protein cytoplasmic dynein, which is essential to power the oscillations (Yamamoto et al., 1999), but has no effect on Mcp5 localization at the cortex (Saito et al., 2006; Yamashita and Yamamoto, 2006). We set up a cross between parental cells containing a deletion of the dynein heavy chain (Dhc1) gene, but containing differently labeled mitochondria, and visualized the distribution of mitochondria in the resulting zygotes and asci. We observed that the parental mitochondria remained segregated in both horsetail zygotes as well and asci (Figures 4A and B, Movie S12 and S13) indicating that the nuclear oscillations had no role to play in the segregation of parental mitochondria.

**Figure 4:**
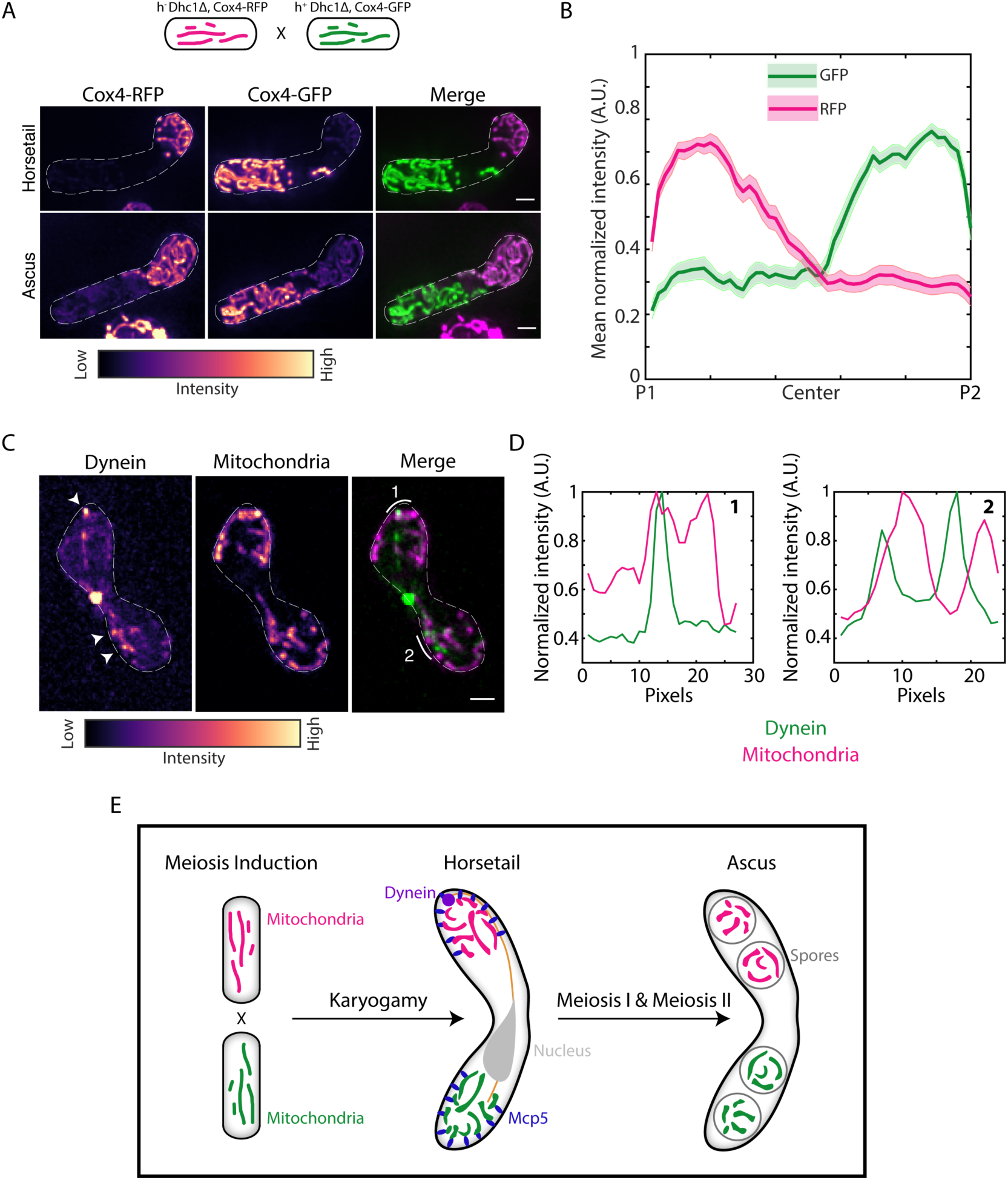
Dynein and mitochondria do not bind to the same Mcp5 foci. **(A)** Schematic of the cross performed (top, strain VA091xVA092, see Table S1), maximum intensity-projected images of mitochondria labeled with Cox4-RFP (left) and mitochondria labeled with Cox4-GFP (center) represented in the intensity map to the bottom of the images, and their merge (right) during the early stage (‘Horsetail’, top) and late stage (‘Ascus’, bottom) of meiosis. **(B)** Plot of mean normalized intensities of RFP (magenta line) and GFP (green line) across the length of the cell from the cross indicated in A (*n*=33). Shaded regions represent SEM. **(C)** Maximum intensity-projected images of dynein (left) and mitochondria (center) represented in the intensity map to the bottom of the images, and their merge (right, strain SV91 stained with MitoTracker Deep Red). White arrowheads point to representative dynein spots on the cortex and line 1 and 2 are drawn along the spots whose intensities are plotted in D. **(D)** Normalized intensity of dynein (green) and mitochondria (magenta) along the arrowheads and lines indicated in C. **(E)** Schematic of uniparental mitochondrial inheritance in fission yeast mediated by the tethering of parental mitochondria to the cortex by the dynein anchor Mcp5. In A and C, scale bars represent 2µm, dashed white lines represent cell outlines.

Since we discovered that the CC domain of Mcp5 not only binds dynein, but also mitochondria, we asked if Mcp5 spots containing dynein clusters were capable of tethering mitochondria. We answered this question by visualizing and measuring the colocalization of dynein-GFP spots and Mitotracker-stained mitochondria (Figure 4C). A typical fission yeast zygote exhibits 1-3 cortical dynein spots that are anchored at Mcp5 foci (Thankachan et al., 2017; Tolic et al., 2009). We observed that only 3 of 23 dynein spots (*n*= 11 zygotes), also localized with mitochondria (Figure 4D, Movie S14), indicating that Mcp5 foci that anchored dynein were typically precluded from tethering mitochondria. This result could explain the slightly better mitochondrial segregation phenotype that we observed in Figure 4A and B, since the lack of dynein in these zygotes made a few more Mcp5 foci available for binding by the mitochondria.

A schematic summarizing these results is depicted in Figure 4E.

## Discussion

Uniparental mitochondrial inheritance is a common feature among several eukaryotes, including unicellular fungi such as *Crytptococcus neoformans* and *Ustilago maydis*. In *C. neoformans*, mitochondria from the MATa parent are selectively passed on to progeny by an as yet unknown degradation mechanism that affects the MATα mitochondria (Yan and Xu, 2003; Yan et al., 2007). In *U. maydis,* the a2 strain, and not a1, contributes all of the mitochondria by employing a mechanism that protects a2 mitochondria from degradation due to the interaction of two genes at the a2 mating type locus, Rga2 and Lga2 (Fedler et al., 2009). In mammalian cells, sperm mitochondria typically enter the oocyte post fertilization, but then undergo selective ubiquitination and proteolysis thereby effecting maternal mitochondrial inheritance in the progeny (Sutovsky et al., 1999, 2000).

Here, we have discovered that the unicellular yeast, *S. pombe* also undergoes uniparental mitochondrial inheritance. The progeny of a meiotic cross are thus homoplasmic for either the h+ or h-parental mitochondria. *S. pombe* achieves uniparental inheritance by employing the anchor protein of dynein, Mcp5, to tether mitochondria to the cortex during meiotic prophase. While this mechanism relies on segregating mitochondria by their anchoring to the cortex, other segregation methods are also possible such as the chloroplast inheritance mechanism in the green alga *Cylindrocystis*, where the two chloroplasts from each parent in the zygote do not mix or divide and are then individually distributed to the four meiotic products (Smith, 1950).

In *S. cerevisiae*, Num1 and Mdm36, which are key components of MECA, serve to anchor mitochondria in the mother cell during mitotic anaphase (Lackner et al., 2013). In contrast to fission yeast, Num1 does not participate in the tethering of mitochondria to the cortex during *S. cerevisiae* meiosis due to the programmed destruction of MECA by Ime2-dependent phosphorylation (Sawyer et al., 2018). In *S. pombe*, the expression profile of Mcp5 peaks during meiotic prophase and drops to zero in the subsequent stages (Saito et al., 2006; Yamashita and Yamamoto, 2006) ensuring that while mitochondria are anchored to the cortex during the earliest stages of meiosis, mitochondria are dissociated from the membrane prior to meiotic divisions I and II.

In budding yeast, Num1 cluster formation requires mitochondrial attachment and the resulting clusters of Num1 are required for dynein anchoring (Kraft and Lackner, 2017; Schmit et al., 2018). In these cells, mitochondria and dynein occupy the same Num1 clusters, with both dynein and mitochondria binding to Num1’s CC domain (Lammers and Markus, 2015). While we too found that Mcp5’s CC domain was required for binding to mitochondria, we observed that distinct populations of Mcp5 clusters are required to anchor dynein and mitochondria. Num1 clusters might accommodate both mitochondria and dynein by making a fraction of molecules in the clusters available for dynein binding after mitochondrial association. In fission yeast, the number of dynein molecules that form a cluster is roughly equal to the number of Mcp5 molecules that make up a focus at the cortex (Ananthanarayanan et al., 2013; Thankachan et al., 2017). Therefore, our results are likely a reflection of the stoichiometry of binding between Mcp5 and dynein, that does not allow for mitochondrial binding to a pre-existing Mcp5-dynein spot.

In conclusion, we report that fission yeast achieves uniparental mitochondrial inheritance by anchoring and thereby segregating parental mitochondria during the earliest stages of meiosis. Future studies will help us understand what the role of uniparental inheritance is in wild-type cells and what the consequence of perturbation of this phenomenon would be, particularly in context of deleterious mtDNA mutations.

## Acknowledgements

We thank A. Jeevannavar and H. Honey for help with construction of strains; High Content Imaging Facility, BSSE, IISc, and P. I. Rajyaguru for the use of the InCell 6000, and Deltavision RT microscopes respectively; P. Delivani, F. Ishikawa, M. Takaine, I. Tolic, P. Tran, and NBRP Japan for yeast strains and constructs; L. A. Chacko for fruitful discussions. The research was supported by the Department of Science and Technology (India)-INSPIRE Faculty Award, the Department of Biotechnology (India) Innovative Young Biotechnologist Award, and the Science and Engineering Research Board (SERB, India) Early Career Research Award awarded to V.A.

## Author contributions

KM and VA carried out all the experiments. VA designed the research, analyzed the data, prepared the figures, and wrote the paper.

## Declaration of interests

The authors declare no competing interests.

