## Supplementary Information for "Cortical tethering of mitochondria by the dynein anchor Mcp5 enables uniparental mitochondrial inheritance during fission yeast meiosis"

### Methods

***Strains and media.*** The fission yeast strains used in the study are listed in Table S1. The fission yeast cells were grown on Yeast Extract (YE) medium or Edinburgh Minimal Medium (EMM) with appropriate supplements (Forsburg and Rhind, 2006).

***Construction of strains.*** Strain VA019 was constructed by crossing strain MTY271 (*h- mCherry-atb2:hphMX6 leu1-32 ura-d18*, see Table S1) with strain FY16887 ($h^{90}$ *leu1-32 (mcp5::ura4+)::GFP-mcp5*, see Table S1) following the Random Spore Analysis (RSA) protocol (Forsburg and Rhind, 2006). Similarly, strain VA066 was constructed by crossing strain PT1651 (*h- cox4-RFP:leu1 ade6-M210 leu1-32 ura4-D18*, see Table S1) with strain FY16839 ($h^{90}$ *leu1-32 ura4-D18 mcp5::ura4+,* see Table S1), strain VA074 was constructed by crossing strain PT1650 (*h+ cox4-GFP:leu1 ade6-M210 leu1-32 ura4-D18*, see Table S1) with strain FY16839 ($h^{90}$ *leu1-32 ura4-D18 mcp5::ura4+,* see Table S1) , strain VA080 was constructed by crossing strain PT2244 (*h+ mmb1Δ:Kanr cox4-GFP:leu2 mCherry-atb2:Hygr ade6-m210 leu1-32 ura4-d18*, see Table S1) with strain L972 (*h- WT,* see Table S1), strain VA086 was constructed by crossing strain PT1651 (*h- cox4-RFP:leu1 ade6-M210 leu1-32 ura4-D18*, see Table S1) with strain FY6871 (*h+ ade6-M210 ura4-D18 leu1,* see Table S1), strain VA091 was constructed by crossing strain PT1650 (*h+ cox4-GFP:leu1 ade6-M210 leu1-32 ura4-D18*, see Table S1) with strain FY21150 (*h- leu1 ura4 dhc1Δ::ura4 (DHC106-1),* see Table S1), and strain VA092 was constructed by crossing strain VA086 (*h+ cox4-RFP:leu1 ade6-M210 ura4-D18*, see Table S1) with strain FY21150 (*h- leu1 ura4 dhc1Δ::ura4 (DHC106-1),* see Table S1).

***Induction of meiosis and preparation of cells for imaging.*** Meiosis was induced in $h^{90}$ strains by suspending a loopful of cells in 100µl of 0.85% NaCl and spotting on to sporulation agar (SPAS) plates. For a cross between $h^{+}$ and $h^{-}$, equal amounts of parental strains were re-suspended in NaCl and spotted onto an SPAS plate. The plate was incubated for ~8h and ~15h at room temperature for $h^{90}$ and $h^{+}/h^{-}$ cross respectively before imaging. For imaging, cells were re-suspended in EMM-N and aspirated onto a 2mg/ml lectin (Sigma-Aldrich, St. Louis, MO, Cat. #L2380) coated 0.17mm glass-bottom dish (SPL, Cat. #100350). Cells were allowed to adhere to the glass bottom for 15-20min. Unattached cells were washed out and cells were imaged in EMM-N.

***MitoTracker Deep Red staining.*** For staining mitochondria in Fig. 4C, meiotic cells were washed once with autoclaved water, and stained with 200nM Mitotracker Deep Red FM (ThermoFisher Scientific, Cat. #M22426) dissolved in EMM-N for 20min. Following this, cells were washed thrice with EMM before imaging.

***DAPI vital staining.*** Staining of mtDNA in live cells was performed using DAPI as described previously (Williamson and Fennell, 1979). Briefly, cells were washed once with water, re-suspended in EMM-N containing 10µg/ml DAPI (Sigma-Aldrich, St. Louis, MO, Cat. #D9542) and allowed to incubate at 30°C for 45min, with shaking at 200rpm. The cells were then washed again with water before proceeding with imaging.

***Microscopy.*** All images are deconvolved and obtained using a Deltavision RT microscope (Applied Precision) with a 100×, oil-immersion 1.4 N.A. objective (Olympus, Japan). Excitation of fluorophores was achieved using InsightSSI (Applied Precision) and corresponding filter selection for excitation and emission of DAPI, GFP, RFP and MitoTracker Deep Red. Z-stacks with 0.2µm-step sizes encompassing the entire cell were captured using a CoolSnapHQ camera (Photometrics), with 2X2 binning. The system was controlled using softWoRx 3.5.1 software (Applied Precision) and the deconvolved images obtained using the built-in setting for each channel. The time-lapse images in Fig. 2C were obtained using the confocal mode in the InCell Analyzer-6000 (GE Healthcare, Buckinghamshire, UK) with 60x/0.7 N.A. objective fitted with an sCMOS 5.5MP camera having an x-y pixel separation of 108nm. For GFP and RFP imaging, 488 and 561nm laser lines and bandpass emission filters 525/20 and 605/52nm respectively were employed. Time-lapses for visualization of mitochondrial dynamics during meiosis were captured with a time interval of 15min.

***Intensity profile measurement.*** The intensity of mitochondria and Mcp5 along the length of cells was obtained in FiJi/ImageJ by measuring the average intensity across a segmented line 25-30 pixels in width drawn along the center of the long axis of the cell.

***Image analysis and plotting.*** Intensity profiles were obtained using FiJi/ImageJ software (Rueden et al., 2017; Schindelin et al., 2012). Analysis was performed using custom functions written in Matlab (Mathworks, Natick, MA). All plots were created using Matlab.

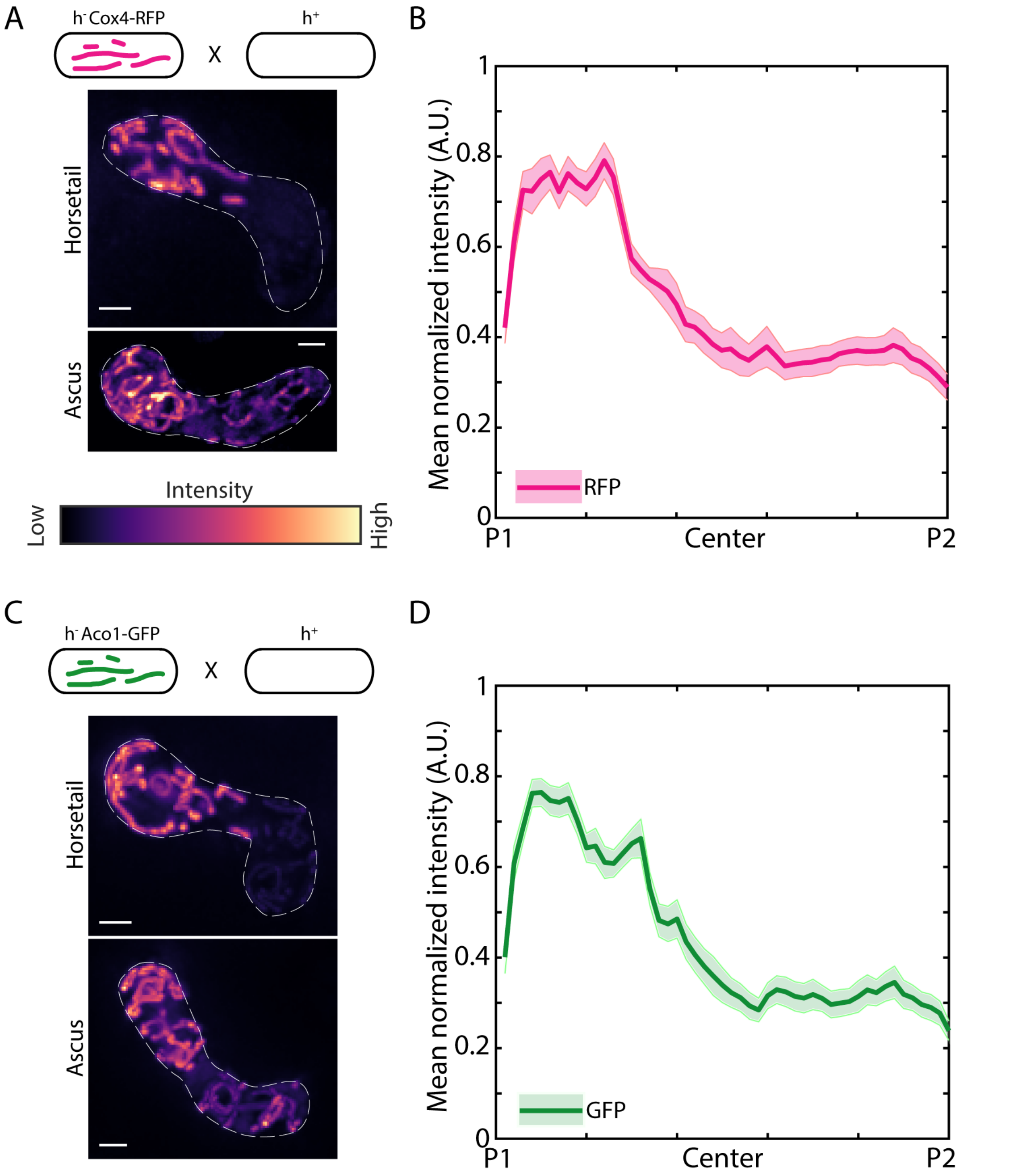


**Figure S1. Parental mitochondria remain segregated during meiosis, Related to Fig. 2. (A)** Schematic of the cross performed (top, strain PT1651xL975, see Table S1), maximum intensity-projected images of mitochondria labeled with Cox4-RFP during the early stage ('Horsetail', top) and late stage ('Ascus', bottom) of meiosis represented in the intensity map to the bottom of the images. **(B)** Plot of mean normalized intensity of RFP (magenta line) across the length of the cell from the cross indicated in A (*n*=13). **(C)** Schematic of the cross performed (top, strain MM3264xL975, see Table S1), maximum intensity-projected images of mitochondria labeled with Aco1-GFP during the early stage ('Horsetail', top) and late stage ('Ascus', bottom) of meiosis represented in the intensity map to the top of the images. **(D)**

Plot of mean normalized intensity of GFP (green line) across the length of the cell from the cross indicated in C (*n*=23). In A and C, scale bars represent 2µm, dashed white lines represent cell outlines. In B and D, shaded regions represent SEM.

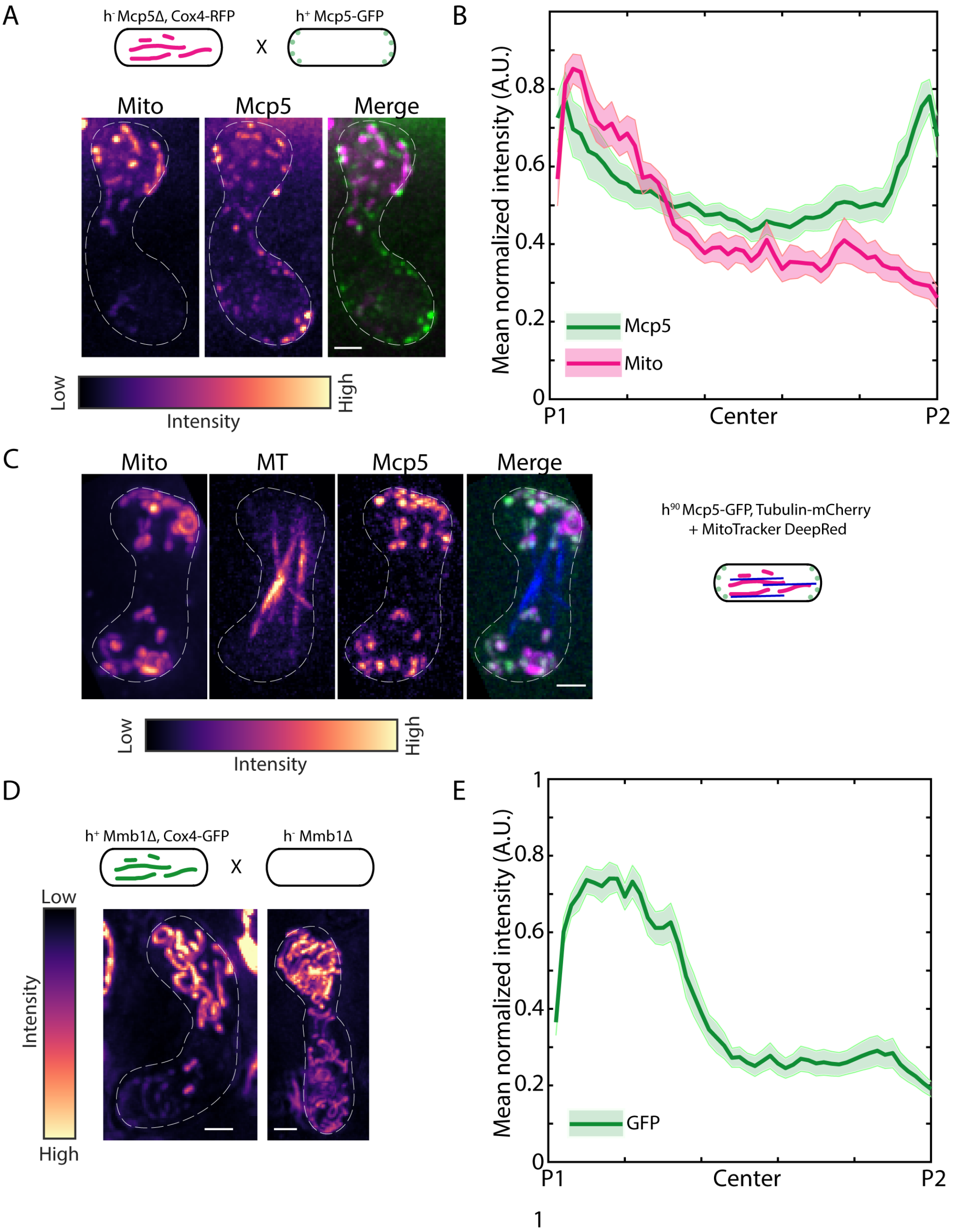


**Figure S2. Mitochondria associate with Mcp5, but not microtubules during meiosis.** (**A**) Schematic of the cross performed (top, strain VA066xFY16854, see Table S1), maximum intensity -projected images of mitochondria labeled with Cox4-RFP (left) and Mcp5 labeled with GFP (center) represented in the intensity map to the bottom of the images, and their merge (right). **(B)** Plot of mean normalized intensities of Mcp5 (green line) and mitochondria (magenta line) across the length of the cell from the cross indicated in A (*n*=10). **(C)** Representative maximum intensity -projected images of mitochondria (first from left), microtubules (second from left) and Mcp5 (second from right) represented in the intensity map to the right of bottom of the images, and their merge (first from right). A schematic of the fluorescence tags is to the right of the images. **(D)** Schematic of the cross performed (top, strain VA080xPT2244, see Table S1), maximum intensity-projected images of mitochondria labeled with Cox4-GFP during the early stage ('Horsetail', top) and late stage ('Ascus', bottom) of meiosis represented in the intensity map to the left of the images. **(E)** Plot of mean normalized intensity of GFP (green line) across the length of the cell from the cross indicated in D (*n*=17). In A, C and D, scale bars represent 2µm, dashed white lines represent cell outlines. In B and E, shaded regions represent SEM.

### Movie captions

**Movie S1.** 3D projection of microtubules (green) and mitochondria (magenta) in a cross between strain KI001 and PT1651 (see Table S1). This movie corresponds to Fig. 1A.

**Movie S2.** 3D projection of nucleus (green) and mitochondria (magenta) in a cross of strain FY15112 (see Table S1). This movie corresponds to Fig. 1B.

**Movie S3.** 3D projection of GFP-labelled mitochondria (green) and RFP-labelled mitochondria (magenta) in a cross between strain PT1650 and PT1651 (see Table S1). This movie corresponds to Fig. 2A 'Horsetail'.

**Movie S4.** 3D projection of GFP-labelled mitochondria (green) and RFP-labelled mitochondria (magenta) in a cross between strain PT1650 and PT1651 (see Table S1). This movie corresponds to Fig. 2A 'Ascus'.

**Movie S5.** Live-cell confocal microscopy of a cross between strain PT1650 and PT1651 (see Table S1). This movie corresponds to Fig. 2C. Scale bar represents 2µm.

**Movie S6.** 3D projection of GFP-labelled mitochondria (green) and DAPI-labelled mtDNA (magenta) in a cross between strain strain PHP14 and PT1650 (see Table S1). This movie corresponds to Fig. 2D.

**Movie S7.** 3D projection of Mcp5 (green) and mitochondria (magenta) in a cross between strain FY16854 and PT1651 (see Table S1). This movie corresponds to Fig. 3A.

**Movie S8.** 3D projection of GFP-labelled mitochondria (green) and RFP-labelled mitochondria (magenta) in a cross between strain VA066 and VA074 (see Table S1). This movie corresponds to Fig. 3C 'Horsetail'.

**Movie S9.** 3D projection of GFP-labelled mitochondria (green) and RFP-labelled mitochondria (magenta) in a cross between strain VA066 and VA074 (see Table S1). This movie corresponds to Fig. 3C ‘Ascus’.

**Movie S10.** 3D projection of mitochondria (warmer colours indicate higher intensities) in a cross between strain FY16897 and VA074 (see Table S1). This movie corresponds to Fig. 3E ‘Horsetail’.

**Movie S11.** 3D projection of mitochondria (warmer colours indicate higher intensities) in a cross between strain FY16897 and VA074 (see Table S1). This movie corresponds to Fig. 3E ‘Ascus’.

**Movie S12.** 3D projection of GFP-labelled mitochondria (green) and RFP-labelled mitochondria (magenta) in a cross between strain VA091 and VA092 (see Table S1). This movie corresponds to Fig. 4A ‘Horsetail’.

**Movie S13.** 3D projection of GFP-labelled mitochondria (green) and RFP-labelled mitochondria (magenta) in a cross between strain VA091 and VA092 (see Table S1). This movie corresponds to Fig. 4A ‘Ascus.

**Movie S14.** 3D projection of dynein (green) and mitochondria (magenta) in a cross of strain SV91 (see Table S1). This movie corresponds to Fig. 4C.

**Table S1. Yeast strains used in this study**

| Name | Genotype | Source |
|---|---|---|
| FY15112 | *$h^{90}$ ade6-216 leu1-32 lys1-131 ura4-D18 hht1::hht1-GFP-HA-Kanr* | NBRP, Japan |
| FY16839 | *$h^{90}$ leu1-32 ura4-D18 mcp5::ura4+* | NBRP, Japan |
| FY16854 | *h+ his2 leu1-32 ura4-D18 mcp5::[mcp5-GFP-3'UTR-Lys3+]* | NBRP, Japan |
| FY16887 | *$h^{90}$ leu1-32 (mcp5::ura4+)::GFP-mcp5* | NBRP, Japan |
| FY16897 | *h- ade6-M216 ura4-D18 (mcp5::ura4+ ::mcp5c-cD* | NBRP, Japan |
| FY6871 | *h+ ade6-M210 ura4-D18 leu1* | NBRP, Japan |
| FY21150 | *h- leu1 ura4 dhc1Δ::ura4 (DHC106-1)* | NBRP, Japan |
| KI001 | *h+ sid4-GFP::kanr kanr -nmtP3-GFP-atb2+ nmt1-pCOX4RFP::leu1+ ura4-D18 ade6-M210* | Iva Tolić, Croatia |
| L972 | *h- WT* | Iva Tolić, Croatia |
| L975 | *h+ WT* | Iva Tolić, Croatia |
| MM3246 | *h- leu1-32 ura4-D18 aco1:GFP:ura4+* | Fuyuki Ishikawa, Japan |
| MTY271 | *h- mCherry-atb2:hphMX6 leu1-32 ura-d18* | Masakatsu Takaine, Japan |
| PHP14 | *h- ade6-M216, leul-32, ptp-1, [$rho^{0}$]* | Thomas Fox, USA |
| PT1650 | *h+ cox4-GFP:leu1 ade6-M210 leu1-32 ura4-D18* | Phong Tran, USA |
| PT1651 | *h- cox4-RFP:leu1 ade6-M210 leu1-32 ura4-D18* | Phong Tran, USA |
| PT2244 | *h+ mmb1Δ:Kanr cox4-GFP:leu2 mCherry-atb2:Hygr ade6-m210 leu1-32 ura4-d18* | Phong Tran, USA |
| SV91 | *$h^{90}$ mcp5-mCherry-kan r dhc1-GFP-Leu2 leu1-32 lys1 ura4-D18* | Iva Tolić, Croatia |
| VA019 | *$h^{90}$ mCherry-atb2:hphMX6 leu1-32 ura-d18 (mcp5::ura4+)::GFP-mcp5* | This study |
| VA066 | *h- cox4-RFP:leu1 mcp5::ura4+ ade6-M210 leu1-32 ura4-D18* | This study |
| VA074 | *h+ cox4-GFP:leu1 mcp5+:: ura4+ ade6-M210 leu1-32 ura4-D18* | This study |
| VA080 | *h- mmb1Δ:Kanr cox4-GFP:leu2 mCherry-atb2:Hygr ade6-m210 ura4-d18* | This study |
| VA086 | *h+ cox4-RFP:leu1 ade6-M210 ura4-D18* | This study |
| VA091 | *h- cox4-GFP:leu1 dhc1Δ::ura4 (DHC106-1) ade6-M210 leu1-32* | This study |
| VA092 | *h+ cox4-RFP:leu1 dhc1Δ::ura4 (DHC106-1) ade6-M210 leu1-32* | This study |
